# DNA methylation age acceleration and risk factors for Alzheimer’s disease

**DOI:** 10.1101/278945

**Authors:** Daniel L McCartney, Anna J Stevenson, Rosie M Walker, Jude Gibson, Stewart W Morris, Archie Campbell, Alison D Murray, Heather C Whalley, David J Porteous, Andrew M McIntosh, Kathryn L Evans, Ian J Deary, Riccardo E Marioni

## Abstract

**INTRODUCTION:** The ‘epigenetic clock’ is a DNA methylation-based estimate of biological age and is correlated with chronological age – the greatest risk factor for Alzheimer’s disease (AD). Genetic and environmental risk factors exist for AD, several of which are potentially modifiable. Here, we assess the relationship associations between the epigenetic clock and AD risk factors.

**METHODS:** Linear mixed modelling was used to assess the relationship between age acceleration (the residual of biological age regressed onto chronological age) and AD risk factors relating to cognitive reserve, lifestyle, disease, and genetics in the Generation Scotland study (n=5,100).

**RESULTS:** We report significant associations between the epigenetic clock and BMI, total:HDL cholesterol ratios, socioeconomic status, and smoking behaviour (Bonferroni-adjusted P<0.05).

**DISCUSSION:** Associations are present between environmental risk factors for AD and age acceleration. Measures to modify such risk factors might improve the risk profile for AD and the rate of biological ageing. Future longitudinal analyses are therefore warranted.

## 1. Introduction

DNA methylation is an epigenetic modification typically characterised by the addition of a methyl group to a ctosine-guanine dinucleotide (CpG). Both genetic and environmental factors influence DNA methylation, which in turn can regulate gene expression [1]. The “epigenetic clock” is an estimation of biological age derived from DNA methylation data, and is strongly correlated with chronological age [2]. From biological age, a measure of age acceleration can be obtained based on the difference between an individual’s biological (estimated) and chronological (actual) age. Age acceleration has been linked to a range of age-related health outcomes, including increased Alzheimer’s disease (AD) pathology [3], reduced cognitive and physical fitness [4], and an increase in all-cause mortality [5]. The epigenetic clock has therefore been proposed as a biomarker of ageing and may be predictive of age-related disorders, such as dementia [6].

Dementia is one of the leading global health concerns of the 21^st^ century. The most common form of dementia is AD. Lifestyle factors such as smoking have been linked to an increased risk of AD [7], as have disease-related factors including type 2 diabetes (T2D) and high blood pressure (HBP) [8, 9]. Moreover, resilience to age-related brain changes (e.g. cognitive reserve), has been linked to AD risk [10]. Factors such as educational attainment and socioeconomic status have been proposed as proxy measures of cognitive reserve, and lower levels of these are established AD risk factors [11, 12]. Genetic studies of AD have revealed several risk factors [13], with the *APOE* locus (encoding apolipoprotein E) being among the strongest [14].

A recent review [15] suggested that up to a third of cases of all-cause dementia might be delayed by actively addressing its modifiable risk factors. In the current study, we investigate the relationship between epigenetic age acceleration and both genetic and environmental AD risk factors in over 5,000 individuals from the Generation Scotland cohort. Two measures of age acceleration were assessed: intrinsic epigenetic age acceleration (IEAA) and extrinsic epigenetic age acceleration (EEAA). These measures are described in greater detail in the methods section. Briefly, IEAA is a measure of age acceleration that is independent of age-related changes in the cellular composition of blood [16], whereas EEAA captures the age-related functional decline of the immune system. Age is the strongest risk factor for AD [17] and epigenetic age is a robust predictor of chronological age. We therefore hypothesise that individuals with poorer profiles for AD risk factors display accelerated ageing in comparison to those with more favourable profiles.

## 2. Methods

### 2.1 The Generation Scotland Cohort

Details of the Generation Scotland Scottish Family Health Study (GS:SFHS) have been described previously [18, 19]. Briefly, the cohort comprises 23,960 individuals, each with at least one family member participating in the study. DNA samples were collected for genotype-and DNA methylation-profiling along with detailed clinical, lifestyle, and sociodemographic data. The current study comprised 5,101 individuals from the cohort for whom DNA methylation data were available. A summary of all variables assessed in this analysis is presented in Table 1.

**McCartney et al. Table 1:**
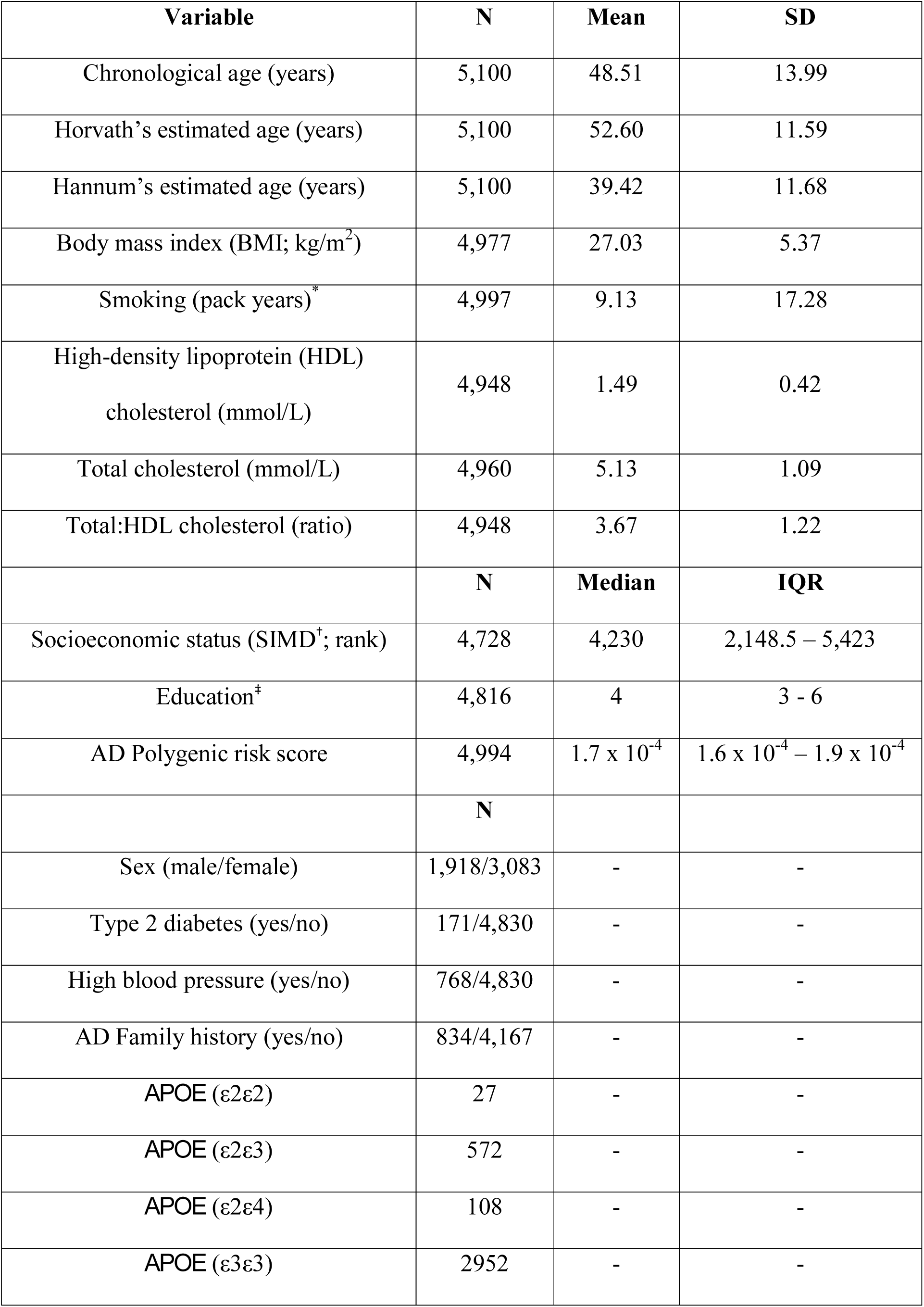
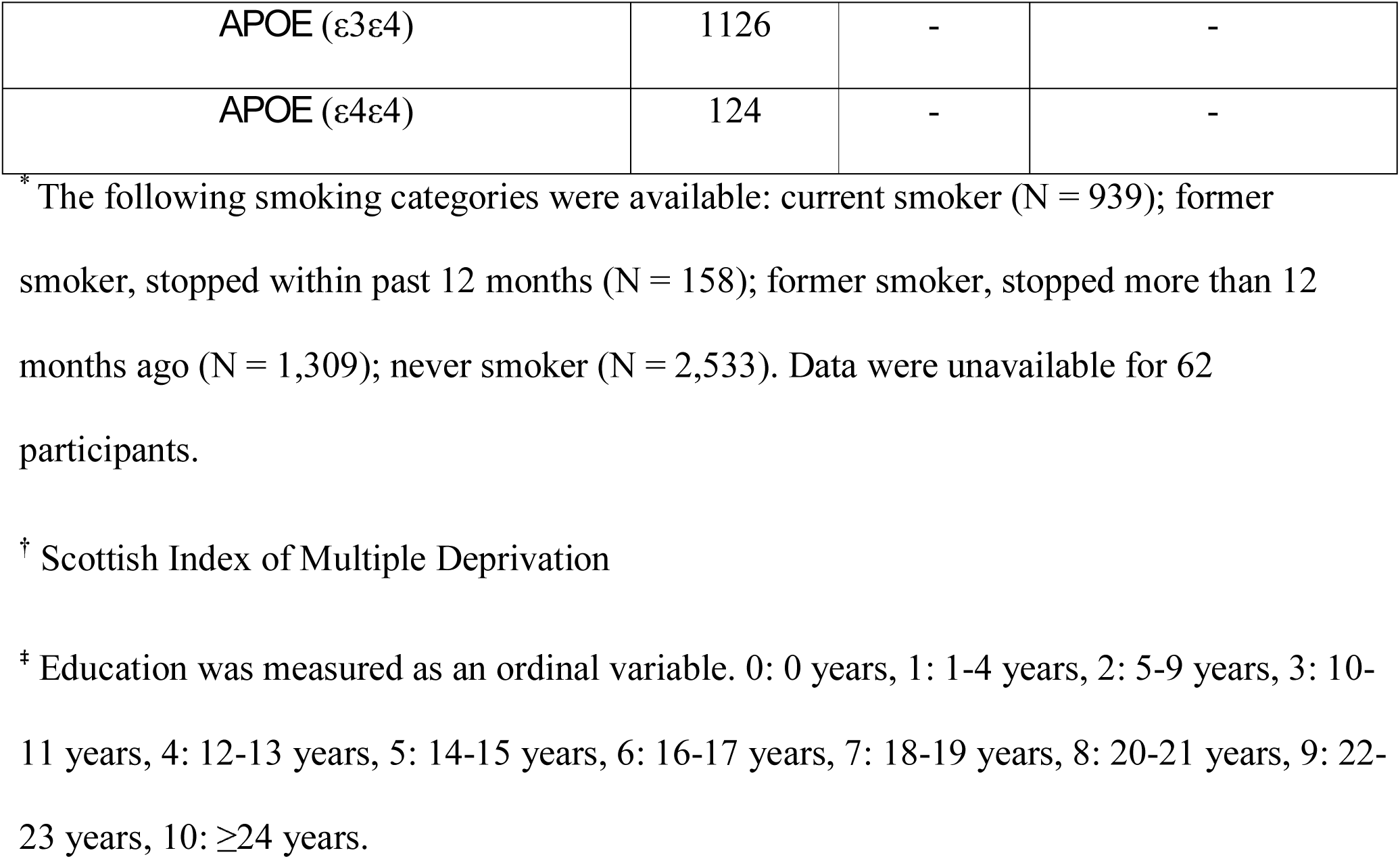
Summary of variables assessed in the Generation Scotland cohort

### 2.2 Ethics

All components of GS:SFHS received ethical approval from the NHS Tayside Committee on Medical Research Ethics (REC Reference Number: 05/S1401/89). GS:SFHS has also been granted Research Tissue Bank status by the Tayside Committee on Medical Research Ethics (REC Reference Number: 10/S1402/20), providing generic ethical approval for a wide range of uses within medical research.

### 2.3 GS:SHFS DNA methylation

Genome-wide DNA methylation was profiled in blood samples from 5,200 individuals using the Illumina HumanMethylationEPIC BeadChips. Quality control was conducted in R [20]. ShinyMethyl [21] was used to plot the log median intensity of methylated versus unmethylated signal per array with outliers being excluded upon visual inspection. WateRmelon [22] was used to remove (1) samples where ≥1% of CpGs had a detection p-value in excess of 0.05 (2) probes with a beadcount of less than 3 in more than 5 samples, and (3) probes where ≥0.5% of samples had a detection p-value in excess of 0.05. ShinyMethyl was used to exclude samples where predicted sex did not match recorded sex. This left a sample of 5,101 available for analysis.

### 2.4 Calculation of age acceleration

Methylation-based estimates of age were calculated using the online age calculator (https://dnamage.genetics.ucla.edu/) developed by Horvath [23]. Normalised GS:SHFS DNA methylation data were used as input for the algorithm and data were underwent a further round of normalisation by the age calculator. Two measures of age acceleration were calculated: intrinsic epigenetic age acceleration (IEAA) and extrinsic epigenetic age acceleration (EEAA). IEAA is defined as the residual term of a multivariate model regressing estimated Horvath methylation age [23] on chronological age, fitting counts of naive CD8+ T cells, exhausted CD8+ T cells, plasmablasts, CD4+ T cells, natural killer cells, monocytes, and granulocytes estimated from the methylation data. IEAA therefore does not consider age-related changes in the cellular composition of blood. Conversely, the estimate of EEAA tracks age-related changes in blood cell composition as well as intrinsic epigenetic changes. EEAA is calculated first by calculating a weighted average of Hannum’s DNA methylation age [24], and three cell types whose abundance is known to change with age (naïve cytotoxic T-cells, exhausted cytotoxic T-cells, and plasmablasts) using the approach described by Klemera and Doubal [25]. EEAA is defined as the residual term of a univariate model regressing the weighted estimated age on chronological age.

### 2.5 Definition of AD risk factors

AD risk factors were divided into four categories: cognitive reserve, disease, lifestyle, and genetics. Cognitive reserve factors comprised education years and socioeconomic status as measured by the Scottish index of multiple deprivation (SIMD). Disease-related factors comprised self-reported type 2 diabetes (T2D) status and high blood pressure (HBP) status. Lifestyle factors comprised smoking pack years (defined as packs smoked per day times years as a smoker), body mass index (BMI), high-density lipoprotein (HDL), total cholesterol and total:HDL cholesterol ratio. Genetic factors comprised family history (defined as having a parent or grandparent with AD), AD polygenic risk score (PGRS), and *APOE* genotype.

### 2.6 Calculation of AD PGRS

PGRS for AD were created for all individuals with genotype data in the GS:SHFS cohort. All autosomal SNPs which passed quality control were included in the calculation of the PGRS for AD (see Supplementary Information for quality control parameters). PGRS for AD were estimated using summary statistics from an independent GWAS of AD (17,008 cases, 37,154 controls), conducted by the International Genomics of Alzheimer’s Project (IGAP) [13]. PGRS were estimated using the PRSice software package, according to previously described protocols [26], with LD threshold and distance threshold for clumping of R^2^ > 0.25 and 250 kb, respectively. After excluding SNPs within a 500kb region of *APOE*, a score was created for each individual, using all possible remaining SNPs, in accordance with previous GS:SFHS analyses [27].

### 2.7 Statistical analysis

Mixed-effects models were performed in R [20], assessing the relationship between epigenetic age acceleration (IEAA and EEAA) and factors related to cognitive reserve, disease, lifestyle, and genetics. Sex and risk factors were fitted as fixed effects and a kinship matrix was fitted as a random effect to control for pedigree structure. Models were built using the *lmekin()* function from the coxme R package [28]. Correction for multiple testing was applied separately to IEAA and EEAA-based analyses using the Bonferroni method.

## 3. Results

### 3.1 Estimation of epigenetic age

Methylation data from 5,101 individuals were submitted to the online age calculator. One individual was flagged for an ambiguous gender prediction and was omitted from downstream analysis, leaving 5,100 individuals. A summary of chronological and estimated ages in the GS:SHFS cohort is provided in Table 1. Both Horvath’s and Hannum’s estimates of biological age were strongly correlated with chronological age (r = 0.94 and 0.93, respectively). As reported previously [29], there was a strong effect of biological sex on age acceleration with men showing greater acceleration than women (Mean EEAA: males = 0.47, females = -0.3, P = 3.58 x 10-12; Mean IEAA: males = 1.13, females = -0.71, P = 8.68 x 10^-53^).

### 3.2 Cognitive reserve and epigenetic age acceleration

Two cognitive reserve factors were evaluated for association with age acceleration: socioeconomic status based on the SIMD, and education years (Table 2; Figure 1). No significant associations were present between these factors and IEAA. Nominally significant negative associations (at P<0.05) were observed between education and SIMD and EEAA (-0.08 years per education category, P = 0.048; -0.26 years per SD increase in SIMD, P = 1.9 x 10^-5^).

**McCartney et al. Table 2:**
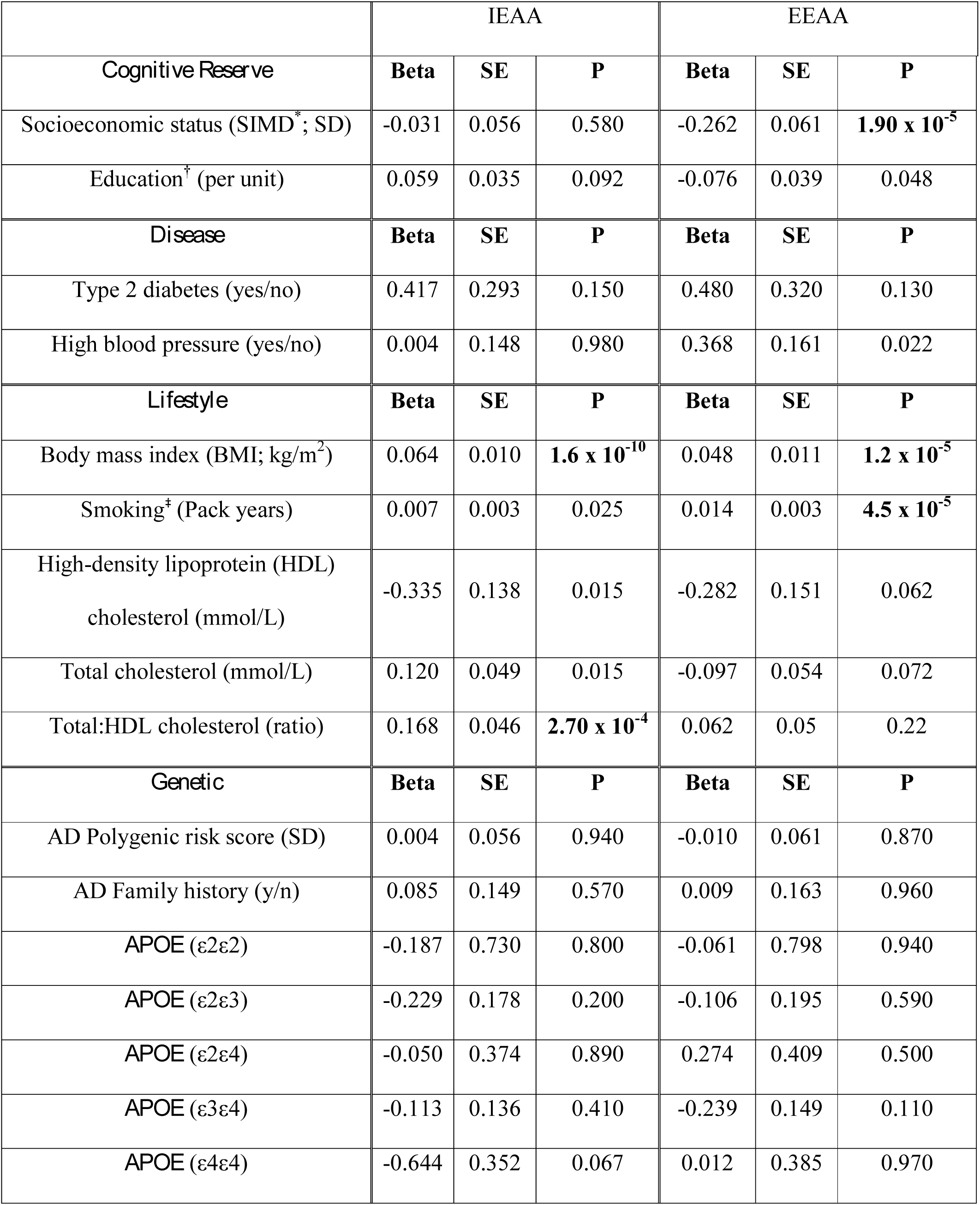
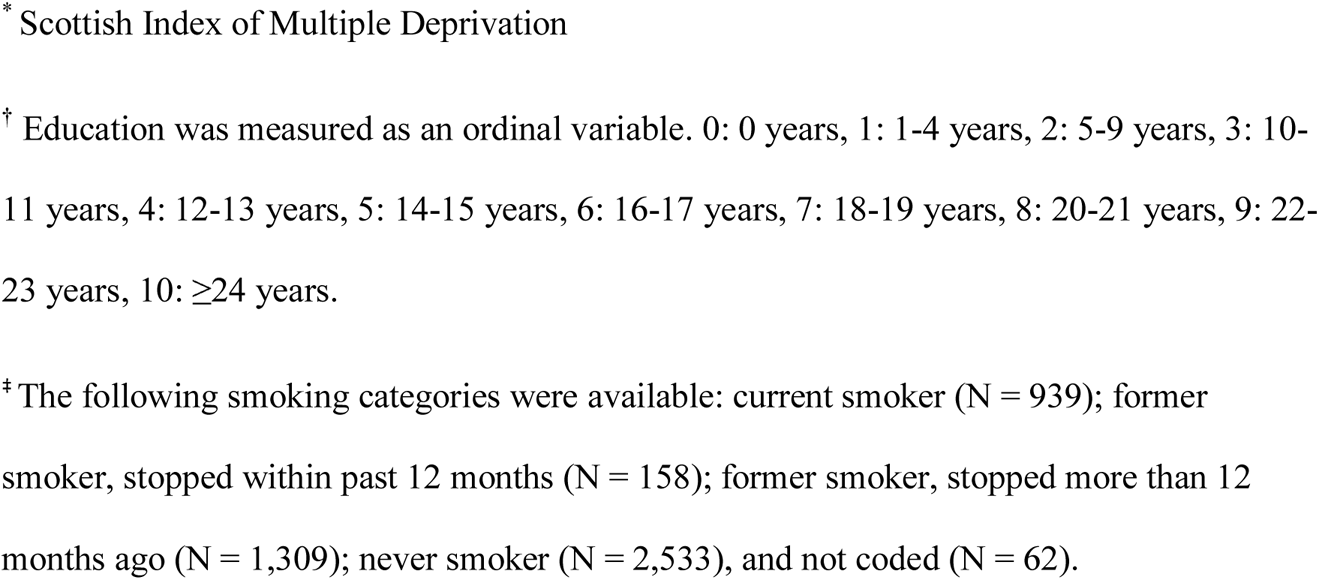
Age acceleration and AD risk factors. Significant associations after accounting for multiple comparisons are highlighted in bold.

|  | IEAA |  |  | EEAA |  |  |
| --- | --- | --- | --- | --- | --- | --- |
| *Cognitive Reserve* | **Beta** | **SE** | **P** | **Beta** | **SE** | **P** |
| Socioeconomic status (SIMD<sup>*</sup>; SD) | -0.031 | 0.056 | 0.580 | -0.262 | 0.061 | **1.90 x 10<sup>-5</sup>** |
| Education<sup>†</sup> (per unit) | 0.059 | 0.035 | 0.092 | -0.076 | 0.039 | 0.048 |
| *Disease* | **Beta** | **SE** | **P** | **Beta** | **SE** | **P** |
| Type 2 diabetes (yes/no) | 0.417 | 0.293 | 0.150 | 0.480 | 0.320 | 0.130 |
| High blood pressure (yes/no) | 0.004 | 0.148 | 0.980 | 0.368 | 0.161 | 0.022 |
| *Lifestyle* | **Beta** | **SE** | **P** | **Beta** | **SE** | **P** |
| Body mass index (BMI; kg/m<sup>2</sup>) | 0.064 | 0.010 | **1.6 x 10<sup>-10</sup>** | 0.048 | 0.011 | **1.2 x 10<sup>-5</sup>** |
| Smoking<sup>‡</sup> (Pack years) | 0.007 | 0.003 | 0.025 | 0.014 | 0.003 | **4.5 x 10<sup>-5</sup>** |
| High-density lipoprotein (HDL) cholesterol (mmol/L) | -0.335 | 0.138 | 0.015 | -0.282 | 0.151 | 0.062 |
| Total cholesterol (mmol/L) | 0.120 | 0.049 | 0.015 | -0.097 | 0.054 | 0.072 |
| Total:HDL cholesterol (ratio) | 0.168 | 0.046 | **2.70 x 10<sup>-4</sup>** | 0.062 | 0.05 | 0.22 |
| *Genetic* | **Beta** | **SE** | **P** | **Beta** | **SE** | **P** |
| AD Polygenic risk score (SD) | 0.004 | 0.056 | 0.940 | -0.010 | 0.061 | 0.870 |
| AD Family history (y/n) | 0.085 | 0.149 | 0.570 | 0.009 | 0.163 | 0.960 |
| *APOE* (ε2ε2) | -0.187 | 0.730 | 0.800 | -0.061 | 0.798 | 0.940 |
| *APOE* (ε2ε3) | -0.229 | 0.178 | 0.200 | -0.106 | 0.195 | 0.590 |
| *APOE* (ε2ε4) | -0.050 | 0.374 | 0.890 | 0.274 | 0.409 | 0.500 |
| *APOE* (ε3ε4) | -0.113 | 0.136 | 0.410 | -0.239 | 0.149 | 0.110 |
| *APOE* (ε4ε4) | -0.644 | 0.352 | 0.067 | 0.012 | 0.385 | 0.970 |

\* Scottish Index of Multiple Deprivation
† Education was measured as an ordinal variable. 0: 0 years, 1: 1-4 years, 2: 5-9 years, 3: 10-11 years, 4: 12-13 years, 5: 14-15 years, 6: 16-17 years, 7: 18-19 years, 8: 20-21 years, 9: 22-23 years, 10: $\geq 24$ years.
‡ The following smoking categories were available: current smoker (N = 939); former smoker, stopped within past 12 months (N = 158); former smoker, stopped more than 12 months ago (N = 1,309); never smoker (N = 2,533), and not coded (N = 62).

### 3.3 Disease-related risk factors and epigenetic age acceleration

We assessed the relationship between age acceleration and two disease-related risk factors: type 2 diabetes and high blood pressure (Table 2; Figure 1). No significant associations were observed between either measure of epigenetic age acceleration and type 2 diabetes. Individuals with high blood pressure displayed an average of 0.37 years of extrinsic age acceleration (P = 0.02).

### 3.4 Lifestyle-related risk factors and epigenetic age acceleration

Four factors related to lifestyle were considered: BMI, smoking habits (pack years), HDL, and total cholesterol (Table 2; Figure 1). Higher values of both measures of epigenetic age acceleration were observed with higher BMI (IEAA: 0.06 years per kg/m^2^ of BMI, P = 1.6 x 10^-10^; EEAA: 0.05 years per kg/m^2^ of BMI, P = 1.2 x 10^-5^), and more pack years (IEAA: 0.007 years per smoking pack year, P = 0.025; EEAA: 0.01 years per smoking pack year, P = 4.5 x 10^-5^). Greater IEAA was associated with lower levels of HDL cholesterol (-0.34 years per mmol/L of HDL. P = 0.015), and higher levels of total cholesterol (0.12 years per mmol/L of total cholesterol, P = 0.015). A significant positive association was present between IEAA and total:HDL cholesterol ratio (beta = 0.168, P = 2.7 x 10^-4^). There were no significant associations observed between EEAA and any of the three cholesterol-related metrics assessed.

### 3.5 Genetic risk factors and epigenetic age acceleration

Three genetic risk factors for AD were assessed for association with age acceleration: family history, AD PGRS, and *APOE* genotype (Table 2; Figure 1). No significant associations were present between any of the genetic risk factors assessed and either measure of epigenetic age acceleration.

### 3.6 Correction for multiple testing

Applying a Bonferroni correction separately for the IEAA and EEAA regressions (P<(0.05/12)=0.0042) identified significant IEAA associations with BMI and total:HDL cholesterol ratio (BMI adjusted P = 2.6 x 10^-9^; total:HDL cholesterol ratio adjusted P = 4.3 x 10^-3^); and significant EEAA associations with SIMD, BMI, and smoking (SIMD adjusted P = 3.0 x 10^-4^; BMI adjusted P = 1.9 x 10^-4^; smoking adjusted P = 7.2 x 10^-4^). Of these, higher age acceleration was associated with higher total:HDL cholesterol ratios, BMI levels, smoking levels and social deprivation.

## 4. Discussion

In the current study, we hypothesised that age acceleration might be associated with dementia risk factors in the Generation Scotland cohort. Using both intrinsic (cell-adjusted) and extrinsic (immune system-associated) estimates of epigenetic age acceleration in a cohort of 5,100 individuals, we identified significant associations between multiple dementia risk factors and age acceleration Several of the AD risk factors associated with age acceleration are potentially modifiable lifestyle factors, suggesting the rate of epigenetic ageing can be altered through behavioural changes.

Biological age has been linked to an increased risk of all-cause mortality and is strongly correlated with chronological age [5]. The epigenetic clock has been proposed as a biomarker of ageing as well as a predictor of an individual’s health and susceptibility to age-related health outcomes [3,5]. As chronological age increases, so too does the risk of dementia. Individuals with greater age acceleration (i.e. with greater epigenetic age relative to chronological age) have slightly poorer cognitive ability [4] and a modest increase in burden of pathological hallmarks of dementia [3].

Of the risk factors assessed, BMI and smoking levels were associated (at a nominal significance threshold) with both estimates of age acceleration. BMI has previously been associated with an increased risk of dementia and AD when high in middle-age and low in old-age [30, 31]. Consistent with our findings, others have observed an association between higher BMI and increased IEAA and EEAA [32]. Previous studies have failed to find associations between smoking levels and epigenetic age acceleration [16, 33]. However, the effect sizes observed in the present study were roughly comparable with those reported by Gao et al. [33] (IEAA: Gao et al. Beta = 0.002-0.0039, current study Beta = 0.007; EEAA: Gao et al. Beta = 0.0073-0.0099, current study Beta = 0.014). Our findings of a significant positive association between self-reported smoking and both measures of age acceleration may be attributable to our larger sample size (N = 4,997 individuals compared to maximum N = 978 individuals with smoking data available [33]) although only EEAA was significantly associated with smoking after correction for multiple testing.

In the present study, factors relating to cholesterol were associated with age acceleration based on the intrinsic (cell-adjusted) estimate of epigenetic age acceleration. HDL levels were negatively correlated with epigenetic age acceleration whereas both total cholesterol levels and total:HDL cholesterol ratio were positively correlated with age acceleration. To our knowledge, significant associations between methylation-based estimates of age acceleration and total:HDL cholesterol ratios have not been reported to date. Consistent with our findings, others have observed an association between lower HDL cholesterol and increased age acceleration [32]. A relationship between increased age acceleration and both total and HDL cholesterol levels using a transcriptomic estimate of biological age has also been reported [34]. HDL cholesterol, colloquially known as “good cholesterol”, primarily functions in lipid transport. Higher levels of HDL cholesterol have been linked to a reduction in cardiovascular disease [35], as well as a decreased risk of AD and dementia [36, 37]. Conflicting evidence exists for the association between mid-life levels of total cholesterol and dementia risk [38, 39]. However, studies have consistently reported an inverse association between total cholesterol levels and AD risk in elderly individuals [40, 41, 42]. Longitudinal analyses have revealed different trajectories of BMI in dementia cases compared to controls [31]. Similarly, longitudinal analyses have also indicated that mid-to-late-life trajectories of cholesterol levels are related to both *APOE* genotype [43] and dementia status [44]. *APOE*, a strong genetic risk factor for AD also functions in lipid transport. The association between cholesterol levels and AD risk, coupled with the functions of *APOE* and other genetic risk factors (e.g. *SORL1*) [13] supports a role for lipid metabolism and transport in dementia [45, 46].

Of the risk factors related to cognitive reserve, both educational attainment and socioeconomic status were associated with EEAA. However, of the two, only socioeconomic status remained significant following Bonferroni correction. Individuals with fewer education years showed increased age acceleration, as did individuals from more deprived socioeconomic backgrounds. Individuals with increased levels of education have displayed delays in the age of onset of dementia [47]. Consistent with our findings, others have reported a similar pattern between EEAA and educational attainment [32, 48]. The biological differences linked to social deprivation are possibly due to the association between socioeconomic status and other, more biologically direct, risk factors for dementia. For example, several lifestyle-related AD risk factors have been shown to be associated with socioeconomic status, including smoking and BMI [49, 50].

No significant associations were observed between either measure of age acceleration and any of the genetic risk factors assessed. Epigenetic age acceleration effects of environmental factors such as smoking and cholesterol may be more visible in blood due to direct contact with the tissue. Whereas genetic risk factors should be consistent across all tissues, it is possible that they only influence epigenetic age acceleration in cell types where AD pathology is primarily observed (i.e. brain tissue).

With a sample size in excess of 5,000 individuals, this is the largest study of DNA methylation-based ageing to date. The use of a comprehensively phenotyped cohort has permitted the assessment of both genetic and environmental AD risk factors and their relationship with epigenetic ageing. This resource is further strengthened by the potential for data linkage to medical records and re-contact of participants, making future longitudinal analyses possible. The cross-sectional design of the current study poses a limitation as it does not permit the assessment of longitudinal changes in age acceleration in response to altered lifestyle habits. However, such a study might be informative in determining whether the trajectory of biological age can be modified through efforts to reduce the risk of AD and other forms of dementia. With the exception of BMI and smoking, significant associations were specific to either IEAA or EEAA. This discordance is possibly due to differences in the two estimates of age acceleration. As described in the methods section, IEAA does not reflect differences in blood cell composition that may be due to age whilst these differences are incorporated into the estimate of EEAA. High blood pressure and both cognitive reserve factors were associated with EEAA, but not IEAA. This may reflect a relationship between these risk factors and immunosenescence. Conversely, the cholesterol-related factors were associated with IEAA but not EEAA, possibly reflecting a relationship between these factors and “pure” epigenetic ageing.

In conclusion, we reported associations between both intrinsic and extrinsic measures of epigenetic age acceleration and environmental AD risk factors. However, no associations were present for the genetic risk factors assessed. At a nominal (P<0.05) significance threshold, IEAA was associated with all of the lifestyle-related factors assessed, whereas EEAA was associated with high blood pressure, BMI, smoking, and both cognitive reserve factors assessed. Following Bonferroni correction, BMI, cholesterol ratios, smoking and socioeconomic status remained significantly associated with epigenetic age acceleration. Risk factors such as cholesterol levels, smoking and BMI can be modulated by behavioural changes with regard to exercise, dietary intake and smoking behaviour. The epigenetic clock is a robust predictor of chronological age, and the greatest risk factor for AD is advanced age [17]. Individuals displaying accelerated ageing have demonstrated increased AD neuropathology and lower cognitive test scores [3, 4]. In the current study, we observed a relationship between age acceleration and AD risk factors. It is reasonable to suggest that, by improving one’s AD risk profile where possible, the process of biological ageing process could be “slowed”.

## Acknowledgements

This work was supported by Alzheimer’s Research UK Major Project Grant [ARUK-PG2017B-10]. Generation Scotland received core funding from the Chief Scientist Office of the Scottish Government Health Directorates [CZD/16/6] and the Scottish Funding Council [HR03006]. We are grateful to all the families who took part, the general practitioners and the Scottish School of Primary Care for their help in recruiting them, and the whole Generation Scotland team, which includes interviewers, computer and laboratory technicians, clerical workers, research scientists, volunteers, managers, receptionists, healthcare assistants and nurses. Genotyping of the GS:SFHS samples was carried out by the Genetics Core Laboratory at the Wellcome Trust Clinical Research Facility, Edinburgh, Scotland and was funded by the Medical Research Council UK and the Wellcome Trust (Wellcome Trust Strategic Award “STratifying Resilience and Depression Longitudinally” (STRADL) [104036/Z/14/Z]. DNA methylation data collection was funded by the Wellcome Trust Strategic Award [10436/Z/14/Z]. The research was conducted in The University of Edinburgh Centre for Cognitive Ageing and Cognitive Epidemiology (CCACE), part of the cross-council Lifelong Health and Wellbeing Initiative [MR/K026992/1]; funding from the Biotechnology and Biological Sciences Research Council (BBSRC) and Medical Research Council (MRC) is gratefully acknowledged. CCACE supports Ian Deary, with some additional support from Dementias Platform UK [MR/L015382/1]. AMM and HCW have received support from the Sackler Institute.

---

**McCartney et al. Figure 1:**
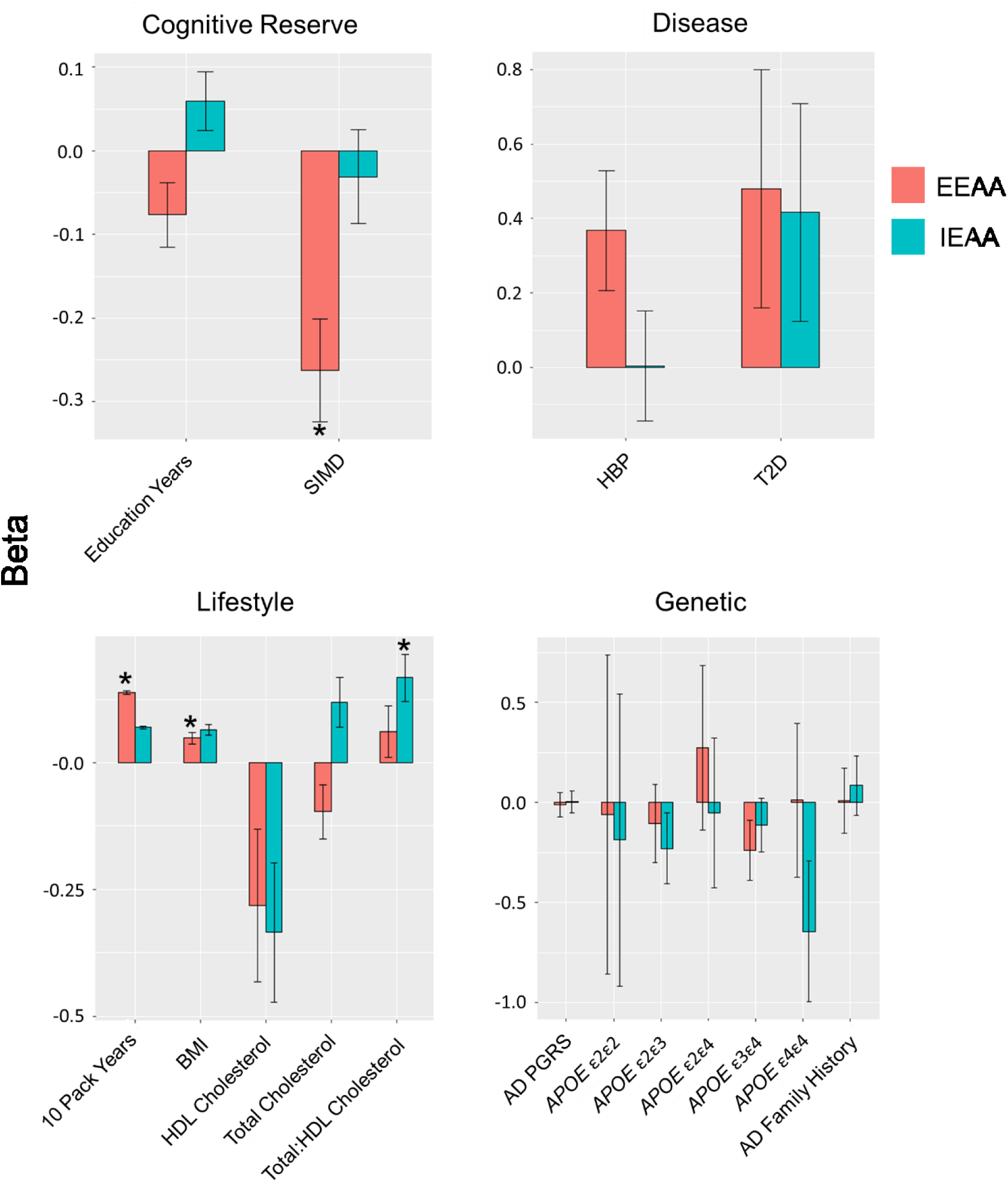
Effects of AD risk factors on age acceleration. Bar plots are separated into four groups of AD risk factors: cognitive reserve, disease, lifestyle, and genetic. Model Beta coefficients (i.e. effect sizes) are presented along the Y-axes while risk factors are presented along the X-axes. Bars are coloured by extrinsic epigenetic age acceleration (EEAA; red) and intrinsic epigenetic age acceleration (IEAA; blue). Error bars show the standard error (SE). Bars accompanied by an asterisk (*) represent measures significantly associated with age acceleration at a Bonferroni P < 0.05. SIMD: Scottish Index of Multiple Deprivation, HBP: High Blood Pressure, T2D: Type 2 Diabetes, BMI: Body Mass Index, HDL: High-Density Lipoprotein cholesterol, AD: Alzheimer’s Disease, PGRS: Polygenic Risk Score. The effect sizes for smoking have been scaled to represent per 10 pack years of exposure. All other effect sizes are per unit increase (disease positive for HBP and T2D and positive family history of AD) with the exception of SIMD and AD polygenic risk score (per SD), and *APOE* status, where the effects are relative to the ε3ε3 reference category.

